# The structural basis for regulation of the nucleo-cytoplasmic distribution of Bag6 by TRC35

**DOI:** 10.1101/154351

**Authors:** Jee-Young Mock, Yue Xu, Yihong Ye, William M. Clemons

## Abstract

The metazoan protein BCL2-associated athanogene cochaperone 6 (Bag6) forms a hetero-trimeric complex with ubiquitin-like 4A (Ubl4A) and transmembrane domain recognition complex 35 (TRC35). This Bag6 complex is involved in tail-anchored protein targeting and various protein quality control pathways in the cytosol as well as regulating transcription and histone methylation in the nucleus. Here we present a crystal structure of Bag6 and its cytoplasmic retention factor TRC35, revealing that TRC35 is remarkably conserved throughout opisthokont lineage except at the C-terminal Bag6-binding groove, which evolved to accommodate a novel metazoan factor Bag6. Remarkably, while TRC35 and its fungal homolog guided entry of tail-anchored protein 4 (Get4) utilize a conserved hydrophobic patch to bind their respective C-terminal binding partners Bag6 and Get5, Bag6 wraps around TRC35 on the opposite face relative to the Get4-5 interface. We further demonstrate that the residues involved in TRC35 binding are not only critical for occluding the Bag6 nuclear localization sequence from karyopherin α binding to retain Bag6 in the cytosol, but also for preventing TRC35 from succumbing to RNF126-mediated ubiquitylation and degradation. The results provide a mechanism for regulation of Bag6 nuclear localization and the functional integrity of the Bag6 complex in the cytosol.

---

Tail-anchored (TA) membrane proteins, which comprise ~2% of eukaryotic genes, contain a single, C-terminal, hydrophobic transmembrane-helix domain (TMD) (1). The TMD, which acts as the endoplasmic reticulum (ER) targeting signal, only emerges from the ribosome after translation termination necessitating post-translational targeting and insertion. Transmembrane domain Recognition Complex (TRC) proteins in mammals and Guided Entry of Tail-anchored proteins (GET) in fungi are the best characterized molecular players in TA targeting (2, 3). In the TRC pathway, TA proteins are captured by smalll-glutamine-rich-tetratricopeptide-repeat protein A (SGTA) after release from the ribosome then transferred to the ATPase TRC40 facilitated by the hetero-trimeric Bag6 complex (which contains Bag6, TRC35, Ubl4A) (4). TA binding stimulates hydrolysis of ATP by TRC40 (5) leading to release of the TRC40-TA complex that then localizes to the ER membrane via its interaction with CAML (6). Release of the TA substrate for insertion is facilitated by calcium modulating ligand (CAML) and tryptophan rich basic protein (WRB) (7). Notably, loss of WRB in mouse cardiomyocytes and hepatocytes results in partial mislocalization of TA proteins (8), suggesting that other pathways can compensate to target TA *in vivo*. Multiple pathways have been implicated in these alternative TA targeting pathways including the signal recognition particle (SRP) (9), SRP-independent targeting (SND) proteins (10), Hsc70 family of chaperones (11), and ubiquilins (12).

Structural information for the mammalian proteins is sparse; however, homology to the equivalent fungal counterparts, Get1-5 and Sgt2, allow for clear parallels. Bag6 is unique among the mammalian proteins, as it appears relatively late in evolution (Fig. S2) and its introduction alters the molecular architecture of the central hand-off complex. For TA targeting, while functionally the heterotrimeric Bag6 complex is equivalent to the fungal heterotetrameric Get4-Get5 complex (4, 13), the additional protein leads to new interactions. In yeast, Get5 (the Ubl4A homolog) has a C-terminal homodimerization domain and an N-terminal heterodimerization domain that forms a complex with Get4 (the TRC35 homolog) resulting in an extended heterotetramer (14, 15).

Crystal structures of fungal Get4 revealed 14 α-helices in an α-solenoid fold that can be divided into an N-terminal and a C-terminal domain (NTD & CTD) (14, 16, 17) (Figs. 1*A* and S1A). The NTD contains conserved residues (Fig. S1C) involved in binding and regulating Get3 (13, 14), while the CTD forms a stable interaction with the Get5 N-terminus (Get5-N) (14, 17). The Get5 interface includes a hydrophobic cleft in Get4 formed by a β-hairpin between α13 and α14 that binds the helix in Get5-N (14). Sequence alignment of Get4/TRC35 and Get5/Ubl4A led to predictions that the Get4 β-hairpin and Get5 N-terminal extension are absent in metazoans (14) (Fig. 1*B*), which suggested a different structural organization for the metazoan Bag6-TRC35-Ubl4A complex.

**Figure 1.**
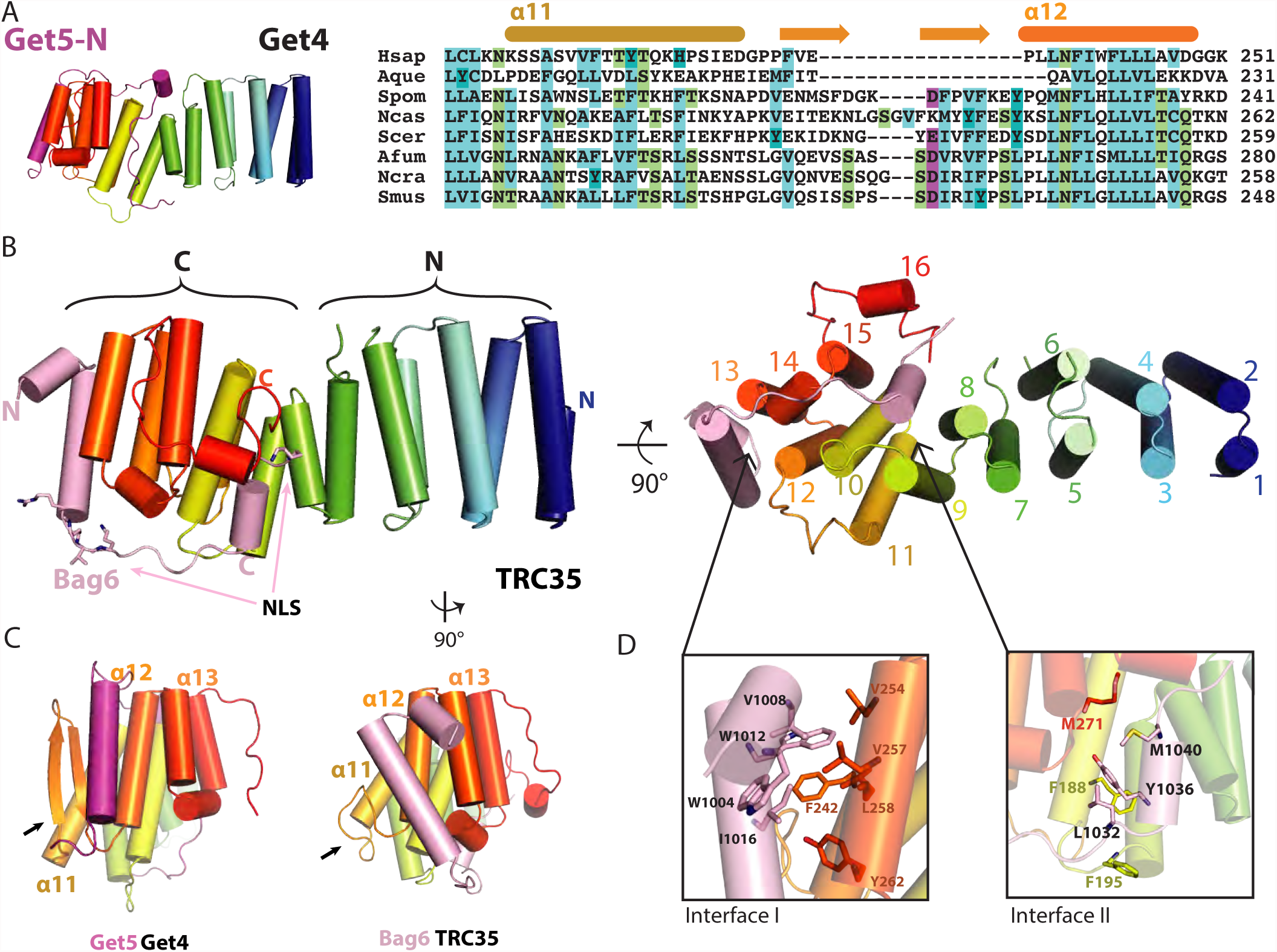
The structure of the Bag6NLS-TRC35 heterodimer. (A) The structure of *Saccharomyces cerevisiae* Get4-Get5N complex (PDBID: 3LKU), Get4 in rainbow and Get5 in magenta. Sequence alignment of TRC35/Get4 from helix 11 to 12 of two metazoan species, *Hsap* (*Homo sapiens*), *Aque* (*Amphimedon queenslandica*), and six fungal species *Spom* (*Schizosaccharomyces pombe*), *Ncas* (*Naumovozyma castelli*), *Scer* (*Saccharomyces cerevisiae*), *Afum* (*Aspergillus fumigatus*), *Ncra* (*Neurospora crassa*), and *Smus* (*Sphaerulina musiva*). The secondary structure based on fungal Get4s is highlighted above the text. (B) Left, the overall structure of Bag6-TRC35 heterodimer in cylinder representation with Bag6 in light pink and TRC35 in rainbow. The nuclear localization sequence is highlighted in sticks on Bag6. Right, a 90° in plane rotated ‘bottom’ view. The 16 helices of TRC35 are labeled from N to C terminus. (C) Comparison of the C-terminal faces of TRC35 and Get4 that bind Bag6 and Get5, respectively. The arrows highlight the significant structural difference in the residues between α11 and α12. (D) Zoomed in view of the regions, defined as interface I and II. Hydrophobic residues that are involved in Bag6-TRC35 dimerization are shown as sticks.

Indeed, in metazoans, Bag6 abrogates the interaction of TRC35 with Ubl4A by separately binding these factors (4, 14, 18) resulting minimally in a heterotrimer. The interface between TRC35 and Bag6 had previously been mapped to the CTD of TRC35 and the region on Bag6 that contains the bipartite nuclear localization sequence (NLS) (4, 19, 20). Overexpression of TRC35 results in Bag6 retention in the cytosol (20), suggesting that TRC35 plays a role in determining the nucleo-cytoplasmic distribution of Bag6. Potential roles for Bag6 in the nucleus are varied and include modulation of p300 acetylation (21-23), facilitation of histone methylation (24, 25), and regulation of DNA damage signaling-mediated cell death (26). How Bag6 localization is regulated remains unclear.

In this study, we present a crystal structure of a complex from humans that includes TRC35 and the TRC35-binding domain of Bag6. The structure of TRC35 reveals the conserved architecture relative to fungal Get4 homologs. Surprisingly, Bag6 utilizes the same conserved pockets on TRC35 that are recognized by Get5 on Get4; yet the extended domain wraps around the opposite face of the α-solenoid. The resulting interaction leads to masking of the bipartite NLS preventing nuclear trafficking of Bag6 by blocking binding to karyopherin α (KPNA), which is demonstrated experimentally. The interaction also prevents RNF126-mediated degradation of TRC35. The combined results suggest a mechanism for regulation of Bag6 nucleo-cytoplasmic distribution.

## The crystal structure of TRC35 – Bag6-NLS complex

A complex between TRC35 (residues 23 to 305), previously shown to be competent for TA transfer (4), and a minimal TRC35-binding domain of Bag6 that contains the NLS (residues 1000-1054) was co-expressed, purified and crystallized. A dataset from a single crystal was collected to 1.8Å resolution and phases were obtained using single wavelength anomalous dispersion from a rubidium derivative. The final model refined to 1.8-40Å had good statistics with an R_work_=15.8% and an R_free_=20.1% and no residues in the disallowed region of the Ramachandran. Full crystallographic statistics are provided in Table S1. All TRC35 residues in the crystallized construct were visible in the electron density except for S137 and G138 that are disordered in the loop between α6 and α7. For Bag6, residues 1002-1043 were resolved with only a few terminal residues ambiguous in the electron density.

TRC35 has the same overall architecture as the fungal Get4 structures (Fig. S1*A*), revealing that the Get4 fold has been conserved across Opisthokonta. Opisthokonta includes metazoan, choanozoan, and fungal lineages (Fig. S2), which share common ancestry as determined by analysis of 16S-like rRNA sequences (27) and several protein sequences (28). The first 15 α-helices form an α-solenoid fold that can be divided into N- and C-terminal halves (Fig. 1*B*). Alignment of the NTD between TRC35 and Get4 (PDB ID: 3LKU) results in an RMSD = 1.380Å for that region, while the equivalent alignment in the CTD results in an RMSD = 3.429Å (Fig. S1*B*). As seen in Get4 (14), there is likely some flexibility between these two halves based on differences across crystal forms. Get4/TRC35 and Get3/TRC40 are conserved throughout eukaryotic evolution and seem to occur as a pair in all opisthokonts, suggesting the essentiality of the two proteins in the pathway (Fig. S2). Consistent with this notion, residues at the TRC35-TRC40 interface (13) and the TRC35-Bag6 interface in TRC35 are highly conserved (Figs. S1*C* and S1*D*). As predicted from sequence alignment analysis, the β-hairpin in Get4 is replaced by a shorter loop in TRC35 (Fig. 1*C*, arrows). In fact, sequence alignment of the predicted Get4 proteins of selected opisthokonts revealed that the β-hairpin is unique to the fungal lineage (Figs. S2 and S3). The C-terminal α16 is flanked by two extended stretches that cover part of Bag6 (Fig. 1*B*).

The Bag6 NLS region wraps around the TRC35-CTD (Fig. 1*B*, light pink) at an interface that resembles the interaction of Get5 with Get4 (Fig. 1*C*). The interaction is stabilized by two hydrophobic interface sites. In interface I, Bag6 α1 and α2 dock into a conserved pocket created by α12 and α13 of TRC35 (Fig. S1*D*) and include W1004, V1008 and W1012 from Bag6 and V254, V257, F242, L258 and Y262 from TRC35 (Fig. 1*D*). In fungal Get4-5, the Get5 N-terminal helix docks into a groove formed by α12, α13, and the β-hairpin of Get4 (Fig. 1*C*). The missing β-hairpin in TRC35 results in the Bag6 helix shifting away from α11 towards α12 and β13 near the bottom of TRC35 (Fig. 1*C*, arrows). The differences in the interface result in changes in the arrangement of α11, α13, and α15 of TRC35 (Fig. 1*C*) relative to those of Get4. Interface II is less extensive involving fewer residues, L1032, Y1036, M1040 of Bag6 and F195, M271 of TRC35 (Fig. 1*D*). While the connecting loops between the two interfaces are well-ordered in both contexts, the Bag6 loop makes fewer contacts to TRC35 than the extensive interactions in the Get5-loop to Get4 (Fig. 1*B*).

The structure reveals how TRC35 masks the Bag6 bipartite NLS (Fig. 1*B*). The conserved first basic cluster of the predicted NLS (K_1024_RVK) is sequestered between interfaces I and II (Fig. 1*B*, sticks). Only the first lysine residue of the second cluster (K_1043_RRK) is resolved in our model (Fig. 1*B*, sticks). The truncated C-terminal residues of TRC35 in our construct are predicted to be disordered (29) and include conserved multiple negative charges (five Glu and three Asp (Fig. S3)) that would be in position to mask the second NLS basic cluster, K1043RRK, through charge-mediated interactions.

## Probing the Bag6-TRC35 Interface

To validate the interfaces observed in the structure, we generated alanine mutants for analysis by yeast 2-hybrid analysis. A fragment of Bag6 (residues 951-1126) was attached to the GAL4 transcription activating domain and full-length TRC35 was attached to the GAL4 DNA-binding domain. The transformation of both vectors were confirmed by cell growth on SC-Ura-Leu plates (Fig. S4A & B). Of the single amino acid substitutions (Figs. 2*A*, S4A & B), only one residue TRC35 (Y262A) at the end of interface I (Fig. 1*D*) disrupted the yeast 2-hybrid interaction (Fig. S4*A*). The Bag6 mutations W1004A and W1012A localize at interface I and Y1036A localizes at interface II. While the individual mutations did not show a phenotype, the combination of the two mutations (W1004A/Y1036A or W1012A/Y1036A) synthetically disrupted the interaction (Fig. 2*A*). For mutants that failed to grow, expression was confirmed by immunoblotting (Fig. S4C). The results confirm that both interfaces are critical for forming a stable complex between Bag6 and TRC35.

**Figure 2.**
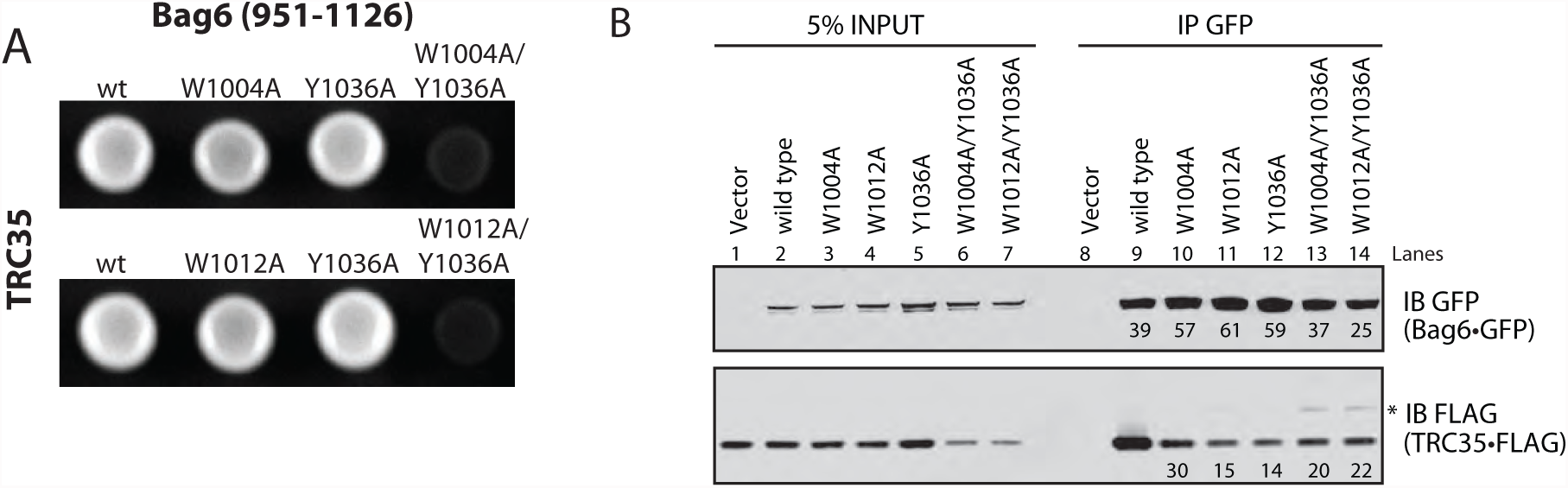
Validation of the Bag6-TRC35 interface. (A) Yeast 2-hybrid assay to validate the interface identified in the crystal structure. Wildtype TRC35 conjugated to the DNA binding domain was expressed with wild-type or mutant Bag6(951-1126) conjugated to the transcription activating domain. Interaction was determined by ability to grow on adenine(-) media. (B) Wild-type or mutant Bag6•GFP was co-expressed in Bag6^-/-^ 293T cells with TRC35•FLAG and immunoprecipitated using anti-GFP antibody. Amount of TRC35 retrieved by Bag6 was assessed by blotting with anti-FLAG antibody. The position of the higher molecular weight TRC35•FLAG is highlighted with an asterisk.

We further validated the interaction of TRC35 with full length Bag6 in the context of a mammalian cellular environment. GFP-tagged wild-type (wt) Bag6 or the mutants Bag6(W1004A), Bag6(W1012), Bag6(Y1036A), Bag6(W1004A/Y1036A), and Bag6(W1012A/Y1036A) were co-expressed with FLAG-tagged TRC35 in CRISPR-mediated Bag6 knock-out 293T cell (30). Bag6 and associated proteins were captured from detergent-derived cell extracts by immunoprecipitating using a GFP-antibody. Viewed by immunoblotting (Figs. 2*B* and S4D & E), each Bag6 mutation resulted in a reduction of the captured TRC35 relative to wtBag6 (compare lane 9 to lanes 10-12). The double mutants (lanes 13 &14) also had reduced capture of TRC35; however, by this assay they unexpectedly appeared to capture more TRC35 than the single mutants. In all the lanes, unexpected higher molecular weight TRC35 bands appeared that became more pronounced in the double mutants (Fig. 2 asterisk and S4D & E). This will be discussed below. Overall, the general effect of mutations at the Bag6 interface are consistent between the yeast 2-hybrid and the mammalian pull-downs.

## Improperly assembled TRC35 is ubiquitylated by RNF126

As noted above, examination of immunoprecipitation results (Figs. 2*B* and S4D & E) revealed two interesting observations: (1) an unexpected increase in TRC35 binding to Bag6 double mutants relative to single mutants (Fig. 2*B* and S4D & E, compare lanes 11-12 vs. lanes 13-14) and (2) the appearance of higher molecular weight products of TRC35 captured by Bag6 double mutants (Figs. 2*B* and S4D & E, asterisk). Evidence in the literature supports that the stability of TRC35 requires forming a proper complex with Bag6 (18, 26). Given the implication of Bag6 as a chaperone holdase in protein quality control processes such as mis-localized protein degradation (31) and ER-associated protein degradation pathways (20, 32), we postulated that TRC35 mutants that fail to form a complex with Bag6 at its physiological binding site are unstable and become a target for Bag6-mediated degradation. In this case, unassembled TRC35 would become a target for Bag6 dependent degradation as a result from TRC35 binding to the Bag6 substrate-binding site. Because the Bag6 construct used in our yeast 2-hybrid studies lacks the substrate binding domain, increased binding is not observed in the double mutant experiments. If this model is correct, the higher molecular weight bands observed in figure 2 and S4D & E should be ubiquitylated TRC35 bound to an alternative substrate-binding site on Bag6.

To verify a role for Bag6 in TRC35 degradation, Bag6 deficient cells were used to co-express TRC35, Bag6 variants and ubiquitin with the expectation that destabilized TRC35 would be ubiquitylated. Cells were transfected with TRC35•FLAG, Bag6•GFP, and HA•ubiquitin. Proteins bound to Bag6 were first immunoprecipitated with GFP antibody (Fig. S5 A & B, lanes 2-7). To remove other ubiquitylated Bag6 substrates (20, 33, 34), the samples obtained from the GFP immunoprecipitation were subject to a second round of immunoprecipitation using anti-FLAG beads under denaturing conditions (Fig. S5A & B, lanes 9-14). Immunoblotting analysis of the samples from the first round of immunoprecipitation showed that all Bag6 variants pulled down ubiquitylated proteins, suggesting that these mutations did not affect its substrate-binding activity (Fig. S5A & B, lanes 2-7). After the second immunoprecipitation with FLAG antibody, TRC35 associated with wtBag6 carried a small amount of ubiquitin conjugates (Fig. S5A & B, lane 9) while those associated with Bag6 variants that disrupted physiological association with TRC35 carried significantly more ubiquitin conjugates (Fig. S5A & B, lanes 10-14). Compared to single Bag6 mutations, TRC35 bound to the double mutants had the highest ubiquitin-modified to unmodified TRC35 ratio (Fig. S5A & B, compare lanes 10-12 to 13-14). Furthermore, the level of TRC35 in cells expressing Bag6 double mutants was consistently lower than in cells expressing wtBag6 (Fig. S5C), which could be rescued by treatment with the proteasome inhibitor MG132 (Fig. 3A lane 2 vs. 4). Treating cells with a proteasome inhibitor also increased ubiquitylated TRC35 that was captured by either wtBag6 or a Bag6 double mutant (Fig. 3A). The accumulation of unmodified TRC35 (Fig. 3A lanes 9 & 10 vs 11 & 12) is probably due to depletion of free ubiquitin in the cell (35). Collectively, these data support the idea that the subset of TRC35 molecules unable to associate with Bag6 via the NLS domain are unstable and become targets for ubiquitin-dependent degradation through a client-chaperone interaction with Bag6. This suggests that the ubiquitylated TRC35 associates with the quality control module (QC) of Bag6 (36).

**Figure 3.**
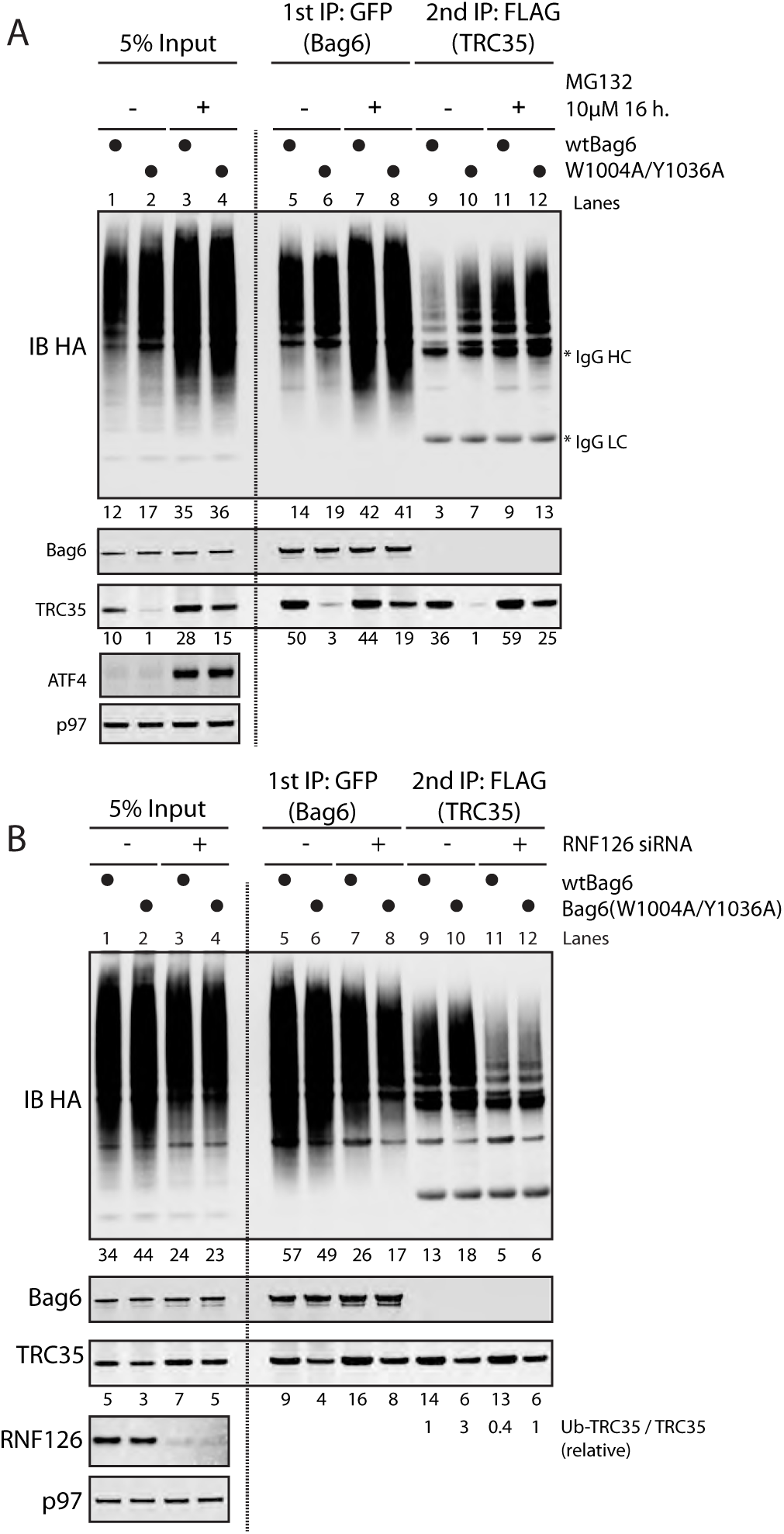
Ubiquitylation of TRC35 in cells expressing Bag6 mutants is dependent on RNF126. (A) Bag6^-/-^ 293T cells co-transfected with plasmids encoding TRC35•FLAG (wt), Bag6•GFP (wt or mutant), and HA•ubiquitin and treated with 10 μM MG132. The cell extracts were subject to two rounds of immunoprecipitation with anti-GFP antibody then with anti-FLAG antibody in denaturing conditions. Ubiquitylation was assessed by immunoblotting with HA antibody. (B) Bag6^-/-^ 293T cells co-transfected with plasmids encoding TRC35•FLAG (wt), Bag6•GFP (wt or mutants), and HA•ubiquitin. The cells were simultaneously treated with 10 μM MG132 and siRNAs against RNF126. The cell extracts were subject to two rounds of denaturing immunoprecipitation with anti-GFP antibody then with anti-FLAG antibody. TRC35 ubiquitylation was assessed by immunoblotting with HA antibody.

To further ensure that unassembled TRC35 is a Bag6-QC substrate, we investigated the effect of the ubiquitin ligase RNF126 on TRC35 ubiquitylation. RNF126 is a Bag6-associated ubiquitin ligase utilized by the Bag6-QC for ubiquitylation of Bag6-associated clients in the cytosol leading to their degradation by the proteasome (36). If TRC35 ubiquitylation is mediated by Bag6-QC, knocking down RNF126 in cells expressing Bag6 mutants would result in reduced ubiquitylation of TRC35, increasing its stability. To test this hypothesis, the effect of RNF126 and proteasome inhibition on TRC35 ubiquitylation and stability were examined.

We examined the effect of siRNA-mediated RNF126 knockdown on TRC35 ubiquitylation. 293T cells co-expressing HA•ubiquitin, TRC35•FLAG and either wt or mutant Bag6•GFP (W1004A/Y1036A or W1012A/Y1036A) were treated with siRNA against RNF126 followed by 6 hour treatment with MG132. Bag6-associated TRC35 was isolated by two rounds of immunoprecipitation. Immunoblotting was used to analyze the relative ratio between ubiquitylated TRC35 and unmodified TRC35 (Ub-TRC35/TRC35). The ratio in cells expressing both wtTRC35 and wtBag6 without RNF126 was defined as 1. In cells expressing Bag6 mutants, ubiquitylation of TRC35 bound to Bag6 is moderately increased (Fig. 3B lane 9 vs. 10); knockdown of RNF126 reduces TRC35 ubiquitylation in cells expressing either wtBag6 or Bag6 double mutants, resulting in ~2-3-fold reduction in Ub-TRC35/TRC35 (Fig. 3B lanes 9 & 10 vs. 11 & 12). Importantly, like MG132 treatment, knockdown of RNF126 rescued the expression of TRC35 in cells expressing the double mutant (Fig. S6), presumably due to increased stability. Together these results demonstrate that ubiquitylation of unassembled TRC35 is modulated by the quality control role of Bag6 and RNF126.

## TRC35 Masks the Bipartite Nuclear Localization Sequence of Bag6 and Retains Bag6 in the Cytosol

To investigate the Bag6-TRC35 interface in the context of Bag6 localization, the mutants identified from yeast 2-hybrid and immunoprecipitation were used for localization studies. Overexpression of TRC35 has been shown to retain Bag6 in the cytosol (20), suggesting that TRC35 binding is required for cytosolic localization of Bag6. Because the mutations identified in our yeast 2-hybrid experiments specifically prevent TRC35 binding at its native binding site that contains the NLS, we postulated that exogenously expressed Bag6 mutants defective in TRC35 binding would localize primarily in the nucleus regardless of TRC35 expression. To test this hypothesis, wt and mutant Bag6 bearing a GFP tag were expressed in HeLa cells with or without TRC35•FLAG and the localization of Bag6 and TRC35 was examined by immunofluorescence.

As expected (20), given the NLS, overexpressed wtBag6 and various Bag6 mutants were localized to the nucleus (Fig. 4*A*). This is likely due to excess Bag6 that cannot be retained in the cytosol by endogenous TRC35. Indeed, when TRC35 is co-expressed with wtBag6, the increased cytosolic pool of TRC35 captures wtBag6 and both proteins stain primarily in the cytosol (Fig. 4*A*). In accordance with the immunoprecipitation results (Fig. 2*B*), introduction of a single mutation—W1004A, W1012A, or Y1036A—results in some Bag6 localization to the nucleus (Fig. 4*B*, *C*, and *D* and table S3). Mutations at both interface I (W1004A or W1012A) and interface II (Y1036A) further reduce the binding affinity between Bag6 and TRC35 (Fig. 2) and Bag6 localizes primarily to the nucleus (Fig. 4*E* & *F* and table S3). These results unequivocally establish TRC35 as a cytoplasmic retention factor of Bag6.

**Figure 4.**
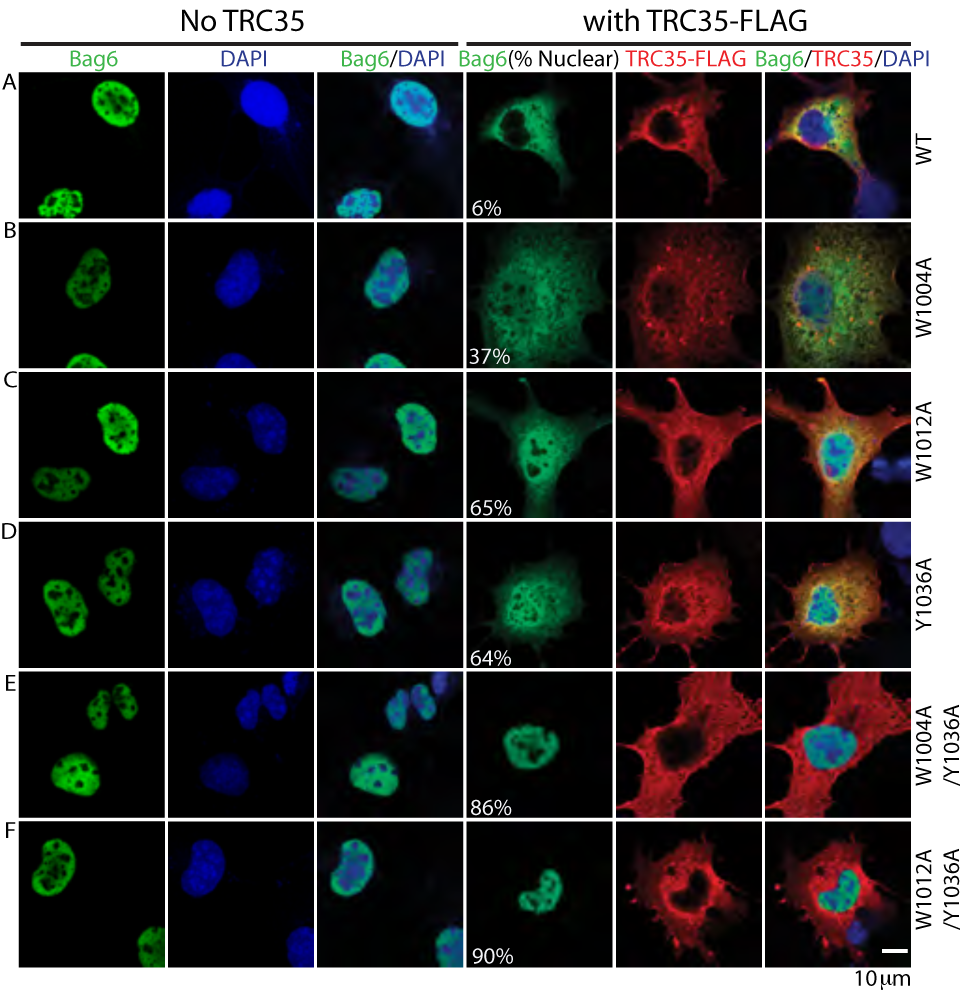
Bag6 mutations at the TRC35 binding site results in nuclear localization of Bag6. (A-F) Representative images of Cos7 cells expressing wtTRC35•FLAG and wt or mutant Bag6•GFP. Cos7 cells were transfected either with Bag6•GFP (wt or mutant) expressing plasmid alone or co-transfected with TRC35•FLAG (wt) expressing plasmid. Cells were stained with anti-GFP (green) and/or anti-FLAG (red) antibodies. DNA was stained with DAPI where indicated (blue). In the Bag6 column, the percent of cells that contain nuclear Bag6 is noted in parenthesis. For each Bag6 variant, approximately 100 cells were counted and categorized as having nuclear or cytosolic Bag6.

## TRC35 has higher affinity for Bag6 than Karyopherin-α 2 (KPNA2)

All molecules destined for the nucleus must move through the nuclear pore complex (NPC), a large assembly that spans the nuclear envelope and facilitates nucleocytoplasmic traffic (37). Macromolecules larger than ~40 kDa cannot freely diffuse through the NPC and require carrier proteins, such as the karyopherin-α (KPNA) and -β families of transport receptors (38, 39). The two basic clusters, R1024KVK and K1043RRK (Fig. 5*A*), in Bag6 are thought to act as a bipartite NLS by specifically recognizing the acidic substrate-binding surface of karyopherins (19). In HeLa cells, the K1045R to S1045L mutation has been shown to abrogate nuclear localization of Bag6 (19). Therefore, TRC35 and KPNA likely bind Bag6 at the fragment that contains the NLS.

**Figure 5.**
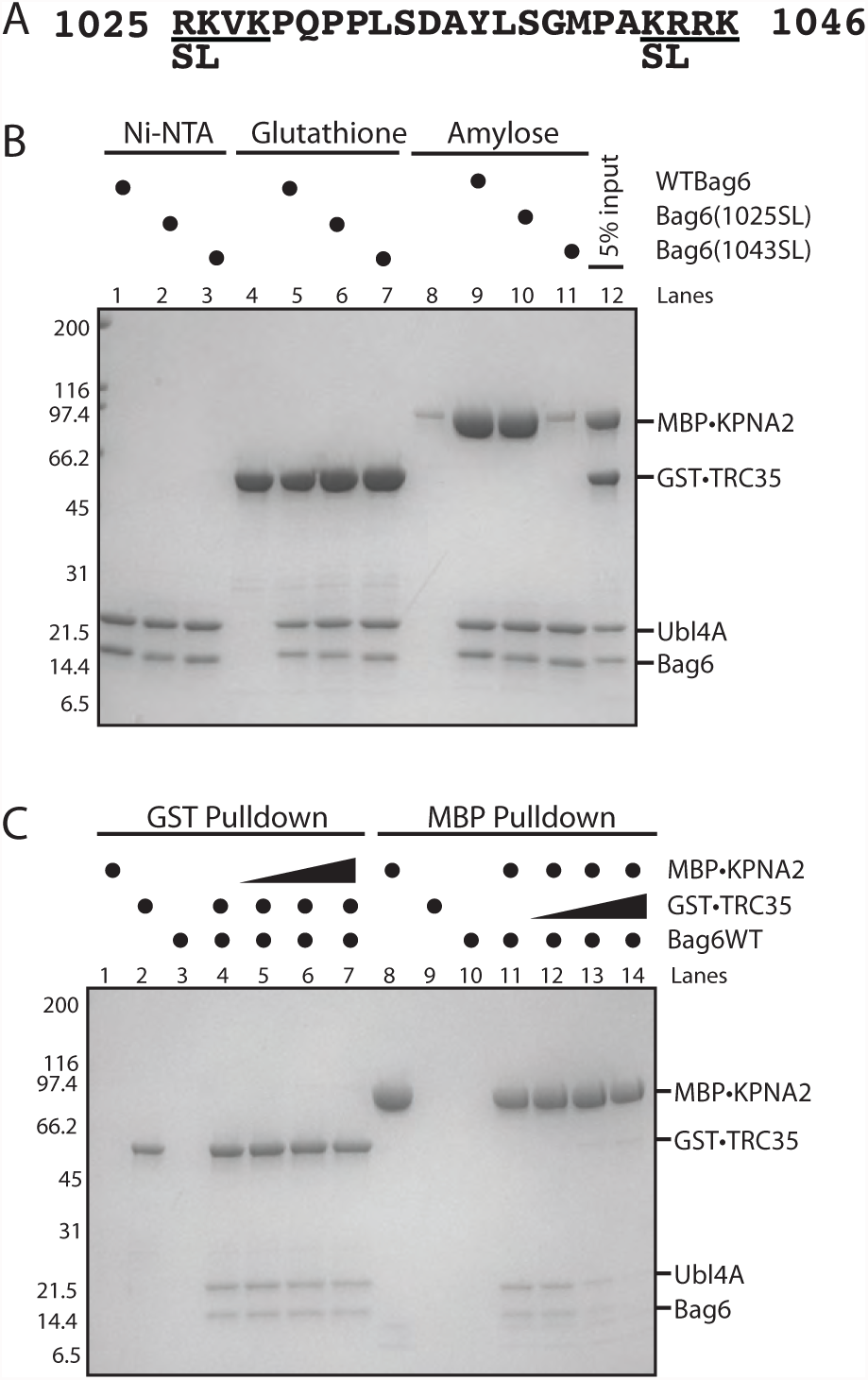
Bag6 is required for TRC35 stability and TRC35 binding precludes karyopherin α binding to Bag6. (A) The putative bipartite nuclear localization sequence of Bag6. The serine and leucine mutations introduced in this study are highlighted. (B) Recombinantly purified Bag6C131-6xHis•Ubl4A (500 pmol) was incubated with two-fold excess GST•TRC35 or MBP•KPNA2 for 20 minutes at room temperature. Ni-NTA beads were used to capture purified Bag6C131-6xHis•Ubl4A and bound factors. (C) Recombinantly purified Bag6C131-6xHis•Ubl4A (500 pmol) was first incubated with either GST•TRC35 or MBP•KPNA2 (500 pmol). The resulting GST•TRC35-Bag6C131-6xHis•Ubl4A complex was incubated with glutathione resin beads, and increasing amounts of MBP•KPNA2 (50 pmol, 250 pmol, or 1000 pmol) was added. The ability of MBP•KPNA2 to displace Bag6C131-6xHis•Ubl4A from GST•TRC35 was examined by eluting the GST•TRC35 and bound Bag6C131-6xHis•Ubl4A from the glutathione resin. The opposite experiment was also carried out using amylose resin beads.

To confirm that the Bag6-NLS is a KPNA binding site and to define the residues involved in KPNA binding, a Bag6 C-terminal 131 residues (Bag6C131) (40) in complex with hexahistidine-tagged full-length Ubl4A (Bag6C131-6xHis•Ubl4A) was purified from *E. coli*. KPNA2 was chosen specifically as it had been seen to interact with Bag6 (41) and could be stably purified. Mutations previously shown to abrogate Bag6 nuclear localization (19) were introduced at either the first basic cluster (1024SL) or the second basic cluster (1045SL) (Fig. 5*A*). The purified Bag6C131-6xHis•Ubl4A variants were incubated with either GST•TRC35 or MBP•KPNA2 lacking importin β binding domain (58-529) and the resulting complexes were isolated using Ni-NTA beads. GST•TRC35 pulled down wild-type, 1024SL, and 1045SL Bag6C131-6xHis•Ubl4A with similar efficiency (Fig. 5*B* lanes 5-7), demonstrating that these residues are not involved in binding TRC35. MBP•KPNA2 formed a complex with Bag6 that can be visualized in a pull-down where MBP•KPNA2 captures wtBag6 (Fig. 5*B* lane 9). Surprisingly, binding was only sensitive to mutation of the second basic cluster which led to significant disruption of the interaction between Bag6C131-6xHis•Ubl4A and MBP•KPNA2 (Fig. 5*B* compare lane 10 and 11). These results show that the residues required for binding KPNA2 are distinct from those required for binding TRC35 despite localizing between key TRC35 binding residues. Moreover, unlike what had been previously suggested (19), the Bag6 NLS is monopartite with the second basic cluster both necessary and sufficient for binding KPNA2. The overlap of the two binding sites provides a simple mechanism for Bag6 retention in the cytoplasm where TRC35 prevents KPNA binding by occlusion.

If binding of TRC35 to Bag6 prevents KPNA-mediated nuclear translocation, only TRC35-free Bag6 should be able to bind KPNA and be translocated to the nucleus. There are several ways in which this could be achieved. One such is that if KPNAs have a higher affinity for Bag6 than TRC35, upregulation of KPNA expression would lead to displacement of TRC35 from Bag6 and formation of a Bag6-KPNA complex. To test this, we compare the relative binding affinities of TRC35 and KPNA2 to Bag6 using an exchange assay. Bag6C131-6xHis•Ubl4A complexes with GST•TRC35 were generated and bound to glutathione affinity resin beads via the GST-tag. After washing, increasing amounts of MBP•KPNA2 were added to the bound beads. The ability of MBP•KPNA2 to displace Bag6C131-6xHis•Ubl4A from GST•TRC35 was determined by the amount of Bag6C131-6xHis•Ubl4A that was eluted from the resin after incubation. In this case, even at the highest concentration tested (2-fold molar excess), there was no significant displacement of Bag6 from TRC35 by KPNA2 (Fig. 5*C* lanes 4-7). Performing the opposite experiment, starting with MBP•KPNA2-Bag6C131-6xHis•Ubl4A on amylose beads, adding excess GST•TRC35 resulted in the dissociation of the MBP•KPNA2-Bag6C131-6xHis•Ubl4A complex (Fig. 5*C* lanes 11-14). These results highlight the stability of the TRC35-Bag6 complex and argue against the ability of KPNA regulation as a means for modulating the nuclear pool of Bag6.

## Discussion

Bag6 is a critical scaffolding factor that forms a stable complex with TRC35 and Ubl4A and is involved in protein targeting, protein quality control, and transcription regulation. While there is ample evidence that Bag6 plays both cytosolic and nuclear roles, it is unclear how the localization of Bag6 is regulated. Here we report the structure of the Bag6 NLS-region bound to its cytosolic retention factor TRC35, and suggest a mechanism for the regulation of Bag6 localization.

The remarkable conservation of TRC35 in both sequence (Fig. S3) and at the structural level (Fig. S1) from yeast to human can be attributed to the role of TRC35 as a hub of protein-protein interactions in the TRC pathway. The most important interaction, based on its complete conservation, is between TRC35 and TRC40 (Fig. S2) and likely drives evolution of components in the pathway. This interaction is critical for the regulation of TA protein transfer (4, 13, 42) and, as shown in fungal Get4, for release of Get3/TRC40 from the ER membrane to restore the cytosolic pool of complex ready for TA protein transfer (43). The importance of linking TRC35 homologs to Ubl4A homologs is sustained in humans despite the loss of structural features that allow these two proteins to interact with each other (14). In this case, the new component Bag6 is introduced, which acts as a scaffold to link TRC35 to Ubl4A. The Bag6-TRC35 structure reveals that TRC35 binds this novel binding partner utilizing conserved patches in a distinct fashion (Fig. 1). This led to the expansion of the TRC35 protein-protein interaction network while maintaining its crucial interaction with TRC40.

Our results also show that TRC35 acts as an intermolecular mask to Bag6 NLS in a role similarly performed in other pathways. Examples of other cytosolic retention factor pairs include IκB and NF-κB (44), HIC and Rev (45) and BRAP2 that retains HMG20A (46). Unlike other cytoplasmic retention factors, which have only been shown to bind their target NLS-containing proteins for occlusion of the NLS, TRC35 also plays a distinct role in the cytoplasmic TA targeting and protein quality control when in complex with Bag6. This dual functionality may be evolutionarily conserved. One study showed that Ubl4A has a nuclear role of promoting STAT3 dephosphorylation (47). This study did not explore the localization of Bag6 relative to Ubl4A, but Ubl4A does not have a NLS and thus is likely to have been translocated with Bag6. In yeast, fungal Get4 binds the N-terminal domain of Get5, which appears to contain a functional NLS that directs Get5 to the nucleus during a ‘Get5-mediated stress response’ (48).

These results allow speculation of possible regulatory mechanism for Bag6 nuclear localization. First, disrupting the Bag6-TRC35 interface by introducing alanine mutations led to ubiquitylation of TRC35 (Fig. S4), which is mediated by RNF126. Knocking down Bag6 has been shown to reduce levels of TRC35 in HeLa cells (26), suggesting that Bag6 is required for TRC35 stability *in vivo*. We also showed that KPNA2 has a lower affinity for Bag6 than TRC35 (Fig. 5*C*), which suggests that for Bag6 to bind KPNA and translocate into the nucleus, it needs to be free of TRC35. The most likely explanation is that cells regulate Bag6 localization by modulating Bag6 levels or decreasing TRC35 levels. In humans, the Bag6 rs3117582 single nucleotide polymorphism found in the promoter region of Bag6 likely affect expression levels and is associated with higher incidence of lung cancer (49-51) and osteoarthritis (52). The localization could also be pre-translationally regulated with differential splicing. In brain and breast tissue, for instance, Bag6 isoforms that lack the NLS are expressed at higher levels than isoforms with the NLS (53).

Our findings establish TRC35 as a universally conserved protein in the opisthokont lineage in both structure and function. These findings support the model in which TRC35 is an important hub of a protein-protein interaction network with its interaction with TRC40 at the core. Higher eukaryotes expanded on this network by the addition of Bag6. Our biochemical studies unequivocally establish TRC35 as a cytosolic retention factor of Bag6 and suggest that Bag6 needs to be in excess for nuclear localization. However, how different cell types regulate Bag6 localization, whether the regulation can be mediated by specific stress, and the biological implications of differential distribution of Bag6 are important questions for future studies.

## Materials and Methods

Detailed descriptions of experiments are provided in SI Materials and Methods. Briefly, for crystallization and *in vitro* binding assays human genes were subcloned and expressed in *E. coli*. Proteins were purified using affinity chromatography, anion-exchange, and size-exclusion chromatography. Truncated fragments of Bag6 and TRC35 were crystallized and the structure was determined using molecular replacement single wavelength anomalous dispersion. The genomes were surveyed for TRC proteins using MEME suite (54). The immunoprecipitation assays were carried out using 293T cell extracts. A combination of three siRNAs against RNF126 was added to 293T cells. 10 μM MG132 was added to inhibit the proteasome. Cos7 cells were used for localization assays.

## Acknowledgements

We thank Daniel Lin and Jens Kaiser for help with data processing. We thank Catherine Day for technical support. We thank members of the laboratory for support and useful discussions. We are grateful to Gordon and Betty Moore for support of the Molecular Observatory at Caltech. We thank the staff at the Stanford Synchrotron Radiation Lightsource for assistance with synchrotron data collection. W.M.C. is supported by NIH grant R01GM097572.

## Author contributions

J.-Y.M. and W.M.C. designed research; J.-Y.M., Y.X., and Y.Y. performed research; Y.X., and Y.Y. contributed new reagents/analytic tools; J.-Y.M., Y.Y., and W.M.C. analyzed data; and J.-Y.M. and W.M.C. wrote the paper.

## The authors declare no conflict of interest

## Data deposition

Coordinates, and structure factors reported in this paper have been deposited in the Research Collaboratory for Structural Bioinformatics Protein Data Bank, www.rcsb.org/pdb/explore.do?structureId=XXX (accession no. XXX).

---

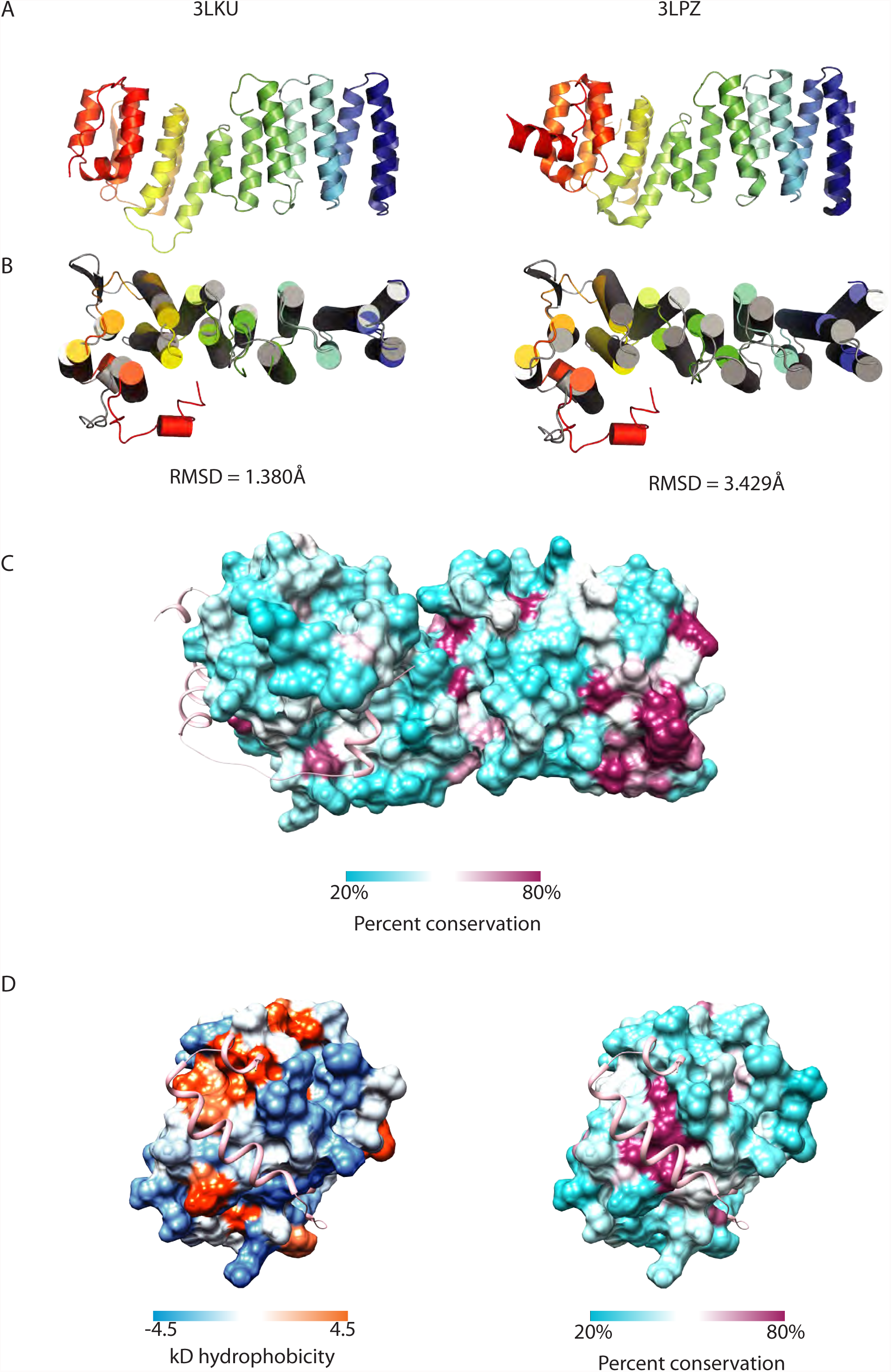
Figure S1

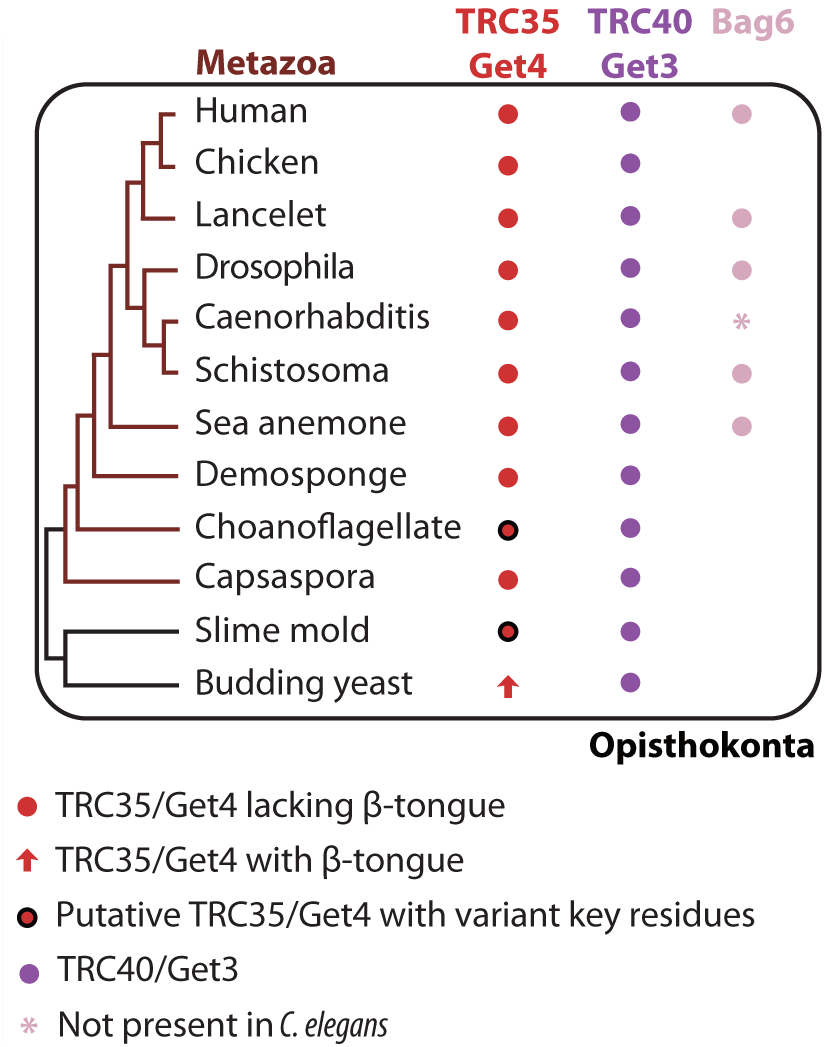
Figure S2

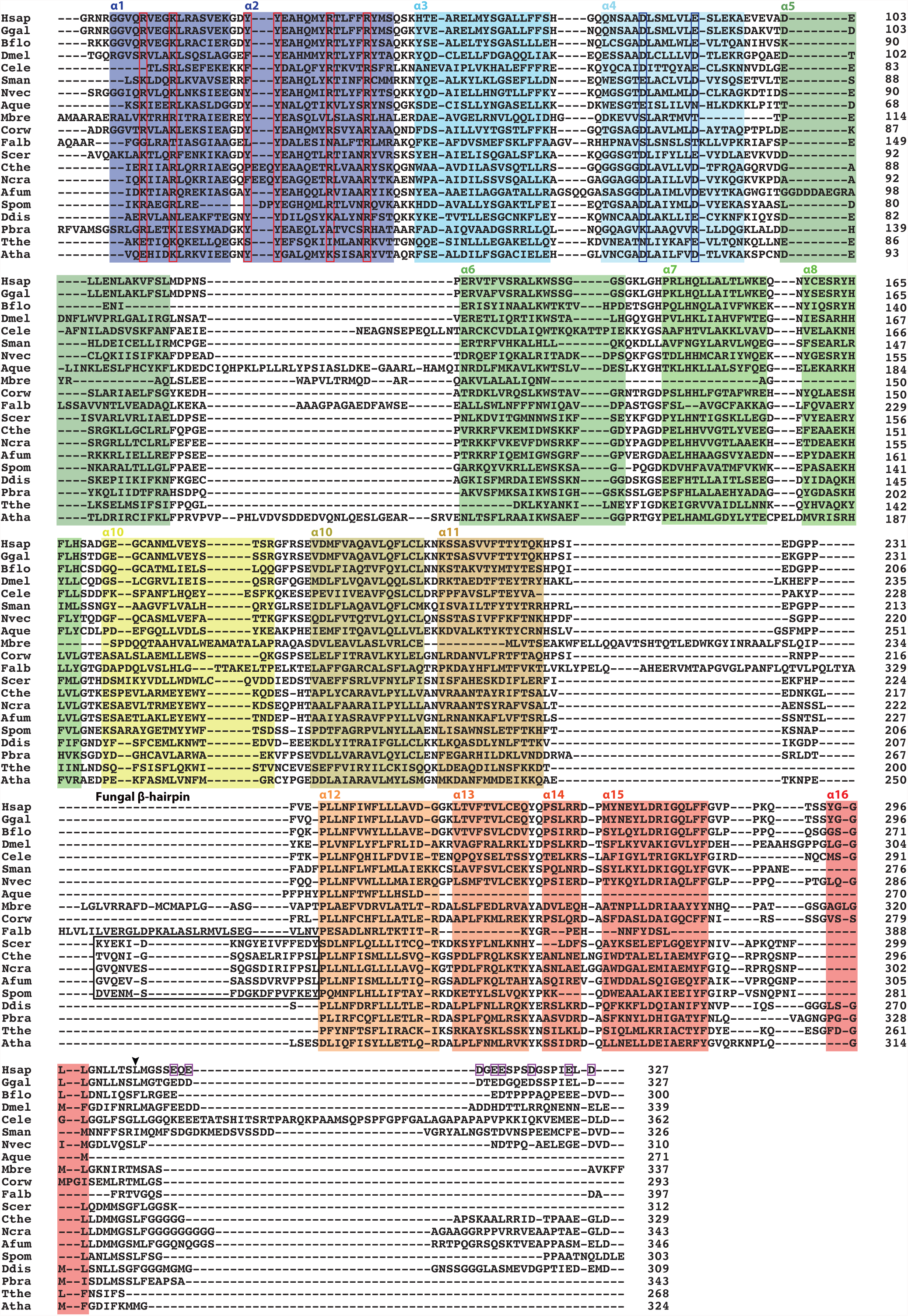
Figure S3

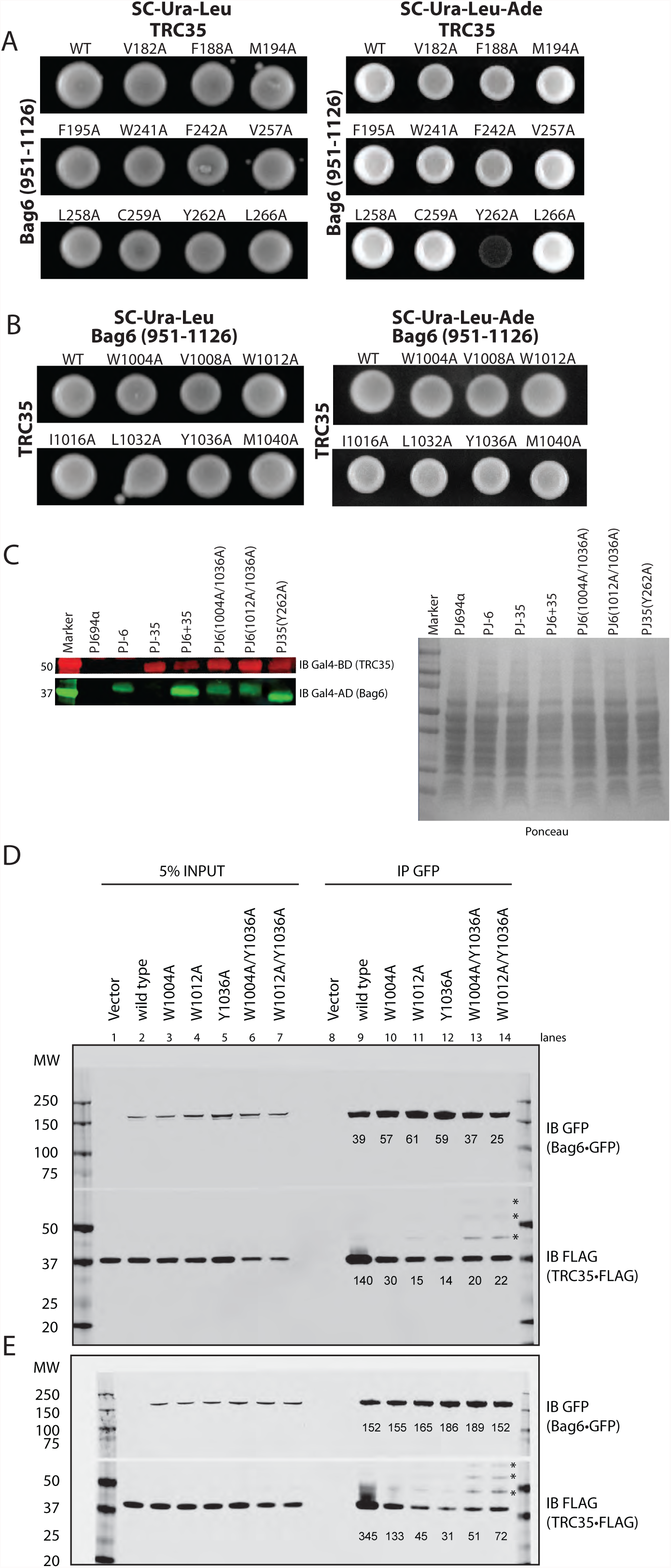
Figure S4

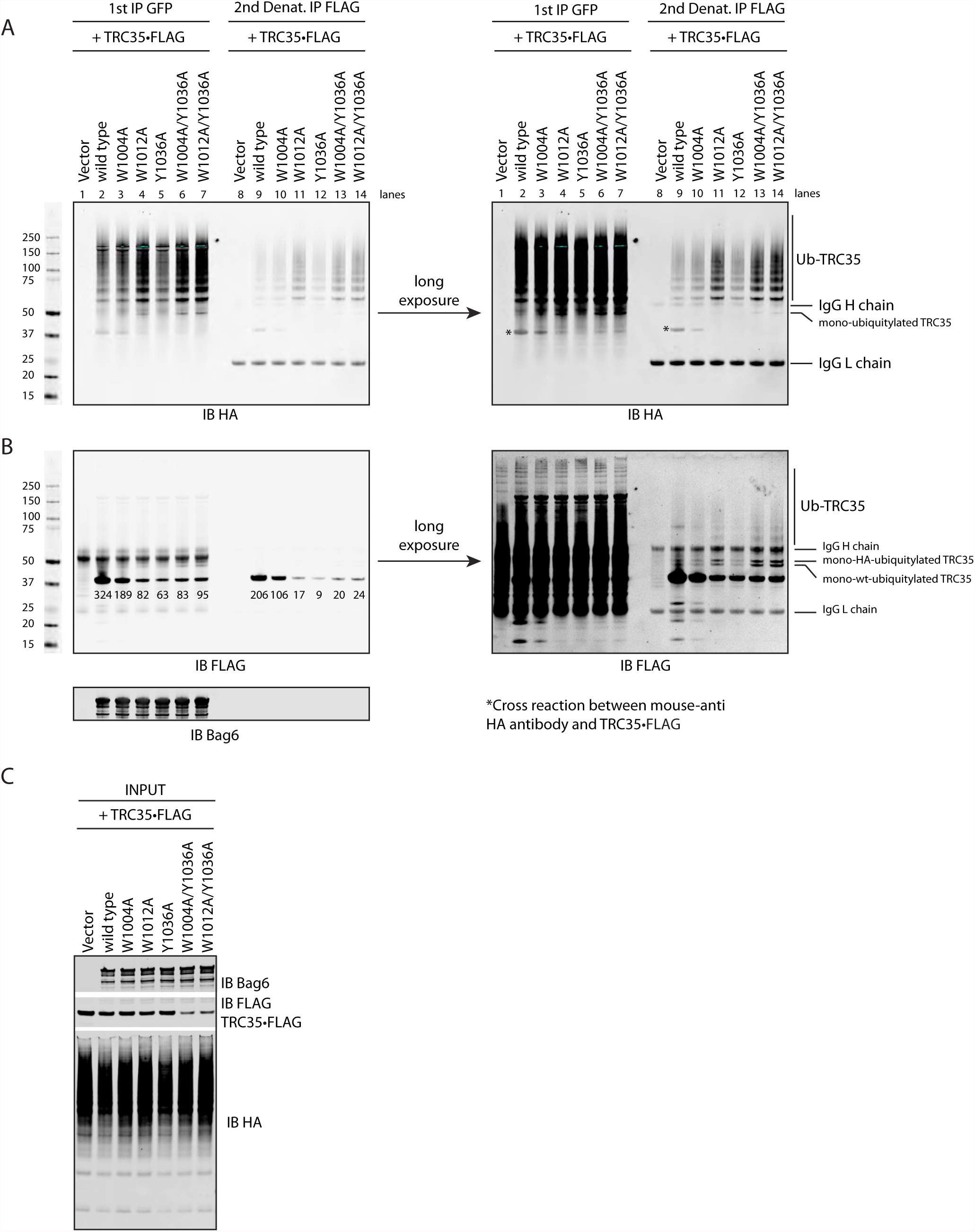
Figure S5

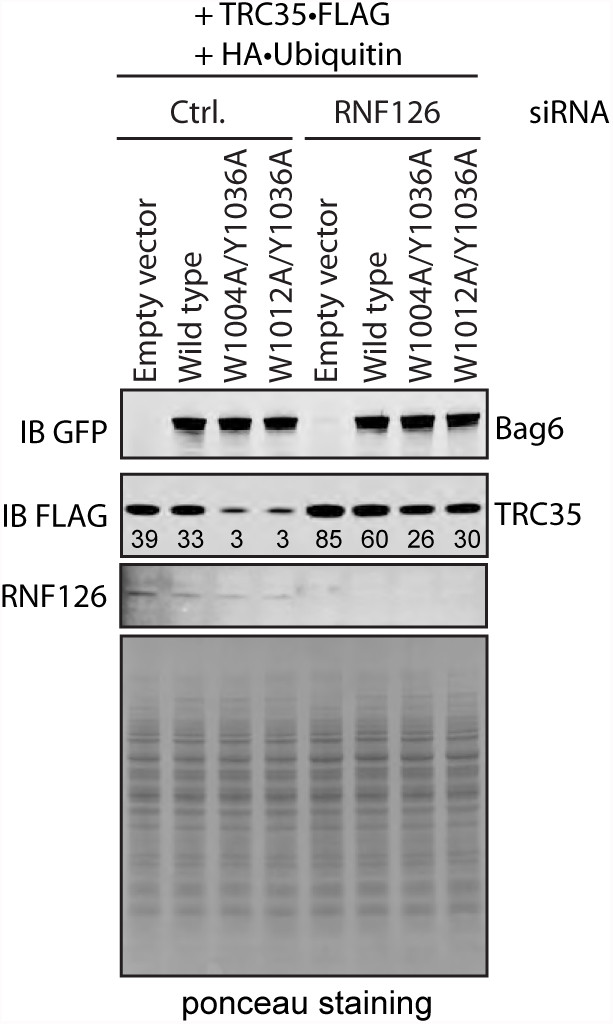
Figure S6

**Table S1.** Crystallographic Analysis

|  | *Native* | *Rubidium* |
| --- | --- | --- |
| **Data collection** |  |  |
| Protein | Bag6-TRC35 | Bag6-TRC35 |
| Synchrotron | SSRL | SSRL |
| Beamline | 12-2 | 12-2 |
| Space group | P21 21 21 | P21 21 21 |
| Cell dimensions |  |  |
| *a*, *b*, *c* (Å) | 41.7, 84.6, 102.6 | 42.0, 84.3, 102.5 |
| $\alpha$, $\beta$, $\gamma$ (°) | 90.0 90.0, 90.0 | 90.0, 90.0, 90.0 |
| Wavelength | 0.9200 | 0.8154 |
| Resolution (Å) | 39.1 – 1.80 (1.92 – 1.80) | 50.0 – 1.99 (2.11 – 1.99) |
| $R_{\text{meas}}$ (%)<sup>a</sup> | 9.3 (64.3) | 16.9 (118.8) |
| $I / \sigma(I)$ | 16.9 (3.8) | 14.9 (2.1) |
| CC<sub>1/2</sub> (%) | 99.9 (95) | 99.9 (88.0) |
| Completeness (%)<sup>a</sup> | 98.7 (92.7) | 99.2 (95.6) |
| No. of observations | 384,047 (60,190) | 655,714 (96,962) |
| No. of unique reflections<sup>a</sup> | 40,776 (6,194) | 47,793 (7,417) |
| Redundancy<sup>a</sup> | 9.4 (9.7) | 13.7 (13.1) |
| **Refinement** |  |  |
| Resolution (Å) | 39.1 – 1.80 |  |
| No. of reflections | 34,334 |  |
| No. of reflections test set | 1,998 |  |
| $R_{\text{work}} / R_{\text{free}}$ | 16.0 / 20.0 | |
| No. atoms (non-hydrogen) | 2725 |  |
| Protein | 2561 |  |
| Water | 152 |  |
| Ligand/Ions | 12 |  |
| *B*-factors |  |  |
| Protein | 31.7 |  |
| Water | 39.0 |  |
| RMSD |  |  |
| Bond lengths (Å) | 0.010 |  |
| Bond angles (°) | 1.16 |  |
| **Ramachandran plot<sup>d</sup>** |  |  |
| Favored (%) | 98 |  |
| Additionally allowed (%) | 2.0 |  |
| Outliers (%) | 0.0 |  |

<sup>a</sup>Highest-resolution shell is shown in parentheses

**Table S2.**

|  | TRC35 | Ubl4A | SGTA | TRC40 | Bag6 |
| --- | --- | --- | --- | --- | --- |
| Homo sapiens | NP_057033.2 | NP_055050.1 | NP_003012.1 | NP_004308.2 | NP_004630.3 |
| Gallus gallus | XP_015149638.1 | XP_015129423.1 | NP_001026550.2 | XP_015158190.1 |  |
| Branchiostoma floridae | XP_002607071.1 | XP_002594764.1 | XP_002591391.1 | XP_002605952.1 | XP_002594881.1 |
| Drosophila melanogaster | NP_649464.1 | NP_650680.1 | NP_609842.1 | NP_610296.2 | NP_648132.2 |
| Caenorhabditis elegans | NP_498034.1 |  | NP_494893.1 | NP_498965.1 |  |
| Schistosoma mansoni | XP_018645653.1 | XP_018645468.1 | XP_018654909.1 | XP_018648676.1 | XP_018648249.1 |
| Nematostella vectensis | XP_001640454.1 | XP_001638929.1 |  | XP_001638254.1 | XP_001635433.1 |
| Amphimedon queenslandica | XP_011404841.1 |  | XP_003387439.1 | XP_003385895.1 |  |
| Monosiga brevicollis | XP_001750773.1 |  | jgi Monbr1 37257 | XP_001743630.1 |  |
| Capsaspora orwazaki | XP_004342754.1 | XP_004347862.1 | XP_004343568.1 | XP_004363354.1 |  |
| Fonticula alba | KCV72828.1 |  | XP_009498174.1 | XP_009492379.1 |  |
| Saccharomyces cerevisiae | NP_014807.3 | NP_014807.3 | NP_014649.1 | NP_010183.1 |  |
| Dictyostelium discoideum | XP_640617.1 | XP_642815.1 | XP_641391.1 | XP_629069.1 |  |
| Trypanosoma brucei |  |  | XP_845518.1 |  |  |
| Plasmodiophora brassicae | CEP00951.1 | CEP02084.1 | CEP01962.1 | CEP03329.1 |  |
| Tetrahymena thermophila | XP_012656464.1 |  |  | XP_001033080.2 |  |
| Arabidopsis thaliana | NP_201127.2 |  | NP_192572.2 | NP_563640.1 |  |

**Table S3.**

|  | Cytosolic Bag6 (# of cells) | Nuclear Bag6 (# of cells) | Total # of cells |
| --- | --- | --- | --- |
| **wtBag6** | 77 | 7 | 84 |
| **Bag6(W1004A)** | 50 | 30 | 80 |
| **Bag6(W1012A)** | 35 | 65 | 100 |
| **Bag6(Y1036A)** | 32 | 56 | 88 |
| **Bag6(W1004A/1036A)** | 18 | 113 | 131 |
| **Bag6(W1012A/1036A)** | 15 | 137 | 152 |

## References

1. Kalbfleisch T, Cambon A, & Wattenberg BW (2007) A bioinformatics approach to identifying tail-anchored proteins in the human genome. Traffic 8(12):1687-1694.

2. Stefanovic S & Hegde RS (2007) Identification of a targeting factor for posttranslational membrane protein insertion into the ER. Cell 128(6):1147-1159.

3. Schuldiner M, et al. (2005) Exploration of the function and organization of the yeast early secretory pathway through an epistatic miniarray profile. Cell 123(3):507-519.

4. Mock JY, et al. (2015) Bag6 complex contains a minimal tail-anchor-targeting module and a mock BAG domain. Proc Natl Acad Sci U S A 112(1):106-111.

5. Rome ME, Rao M, Clemons WM, & Shan SO (2013) Precise timing of ATPase activation drives targeting of tail-anchored proteins. Proc Natl Acad Sci U S A 110(19):7666-7671.

6. Yamamoto Y & Sakisaka T (2012) Molecular machinery for insertion of tail-anchored membrane proteins into the endoplasmic reticulum membrane in mammalian cells. Mol Cell 48(3):387-397.

7. Vilardi F, Lorenz H, & Dobberstein B (2011) WRB is the receptor for TRC40/Asna1-mediated insertion of tail-anchored proteins into the ER membrane. J Cell Sci 124(Pt 8):1301-1307.

8. Rivera-Monroy J, et al. (2016) Mice lacking WRB reveal differential biogenesis requirements of tail-anchored proteins in vivo. Sci Rep 6:39464.

9. Abell BM, Pool MR, Schlenker O, Sinning I, & High S (2004) Signal recognition particle mediates post-translational targeting in eukaryotes. EMBO J 23(14):2755-2764.

10. Aviram N, et al. (2016) The SND proteins constitute an alternative targeting route to the endoplasmic reticulum. Nature 540(7631):134-138.

11. Abell BM, Rabu C, Leznicki P, Young JC, & High S (2007) Post-translational integration of tail-anchored proteins is facilitated by defined molecular chaperones. J Cell Sci 120(Pt 10):1743-1751.

12. Itakura E, et al. (2016) Ubiquilins Chaperone and Triage Mitochondrial Membrane Proteins for Degradation. Mol Cell 63(1):21-33.

13. Gristick HB, et al. (2014) Crystal structure of ATP-bound Get3-Get4-Get5 complex reveals regulation of Get3 by Get4. Nat Struct Mol Biol 21(5):437-442.

14. Chartron JW, Suloway CJ, Zaslaver M, & Clemons WM, Jr. (2010) Structural characterization of the Get4/Get5 complex and its interaction with Get3. Proc Natl Acad Sci U S A 107(27):12127-12132.

15. Chartron JW, VanderVelde DG, Rao M, & Clemons WM, Jr. (2012) Get5 carboxyl-terminal domain is a novel dimerization motif that tethers an extended Get4/Get5 complex. J Biol Chem 287(11):8310-8317.

16. Bozkurt G, et al. (2010) The structure of Get4 reveals an alpha-solenoid fold adapted for multiple interactions in tail-anchored protein biogenesis. FEBS letters 584(8):1509-1514.

17. Chang YW, et al. (2010) Crystal structure of Get4-Get5 complex and its interactions with Sgt2, Get3, and Ydj1. J Biol Chem 285(13):9962-9970.

18. Mariappan M, et al. (2010) A ribosome-associating factor chaperones tail-anchored membrane proteins. Nature 466(7310):1120-1124.

19. Manchen ST & Hubberstey AV (2001) Human Scythe contains a functional nuclear localization sequence and remains in the nucleus during staurosporine-induced apoptosis. Biochem Biophys Res Commun 287(5):1075-1082.

20. Wang Q, et al. (2011) A ubiquitin ligase-associated chaperone holdase maintains polypeptides in soluble states for proteasome degradation. Mol Cell 42(6):758-770.

21. Sebti S, et al. (2014) BAT3 modulates p300-dependent acetylation of p53 and autophagy-related protein 7 (ATG7) during autophagy. Proc Natl Acad Sci U S A 111(11):4115-4120.

22. Sebti S, et al. (2014) BAG6/BAT3 modulates autophagy by affecting EP300/p300 intracellular localization. Autophagy 10(7):1341-1342.

23. Sasaki T, et al. (2007) HLA-B-associated transcript 3 (Bat3)/Scythe is essential for p300- mediated acetylation of p53. Genes Dev 21(7):848-861.

24. Wakeman TP, Wang Q, Feng J, & Wang XF (2012) Bat3 facilitates H3K79 dimethylation by DOT1L and promotes DNA damage-induced 53BP1 foci at G1/G2 cell-cycle phases. EMBO J 31(9):2169-2181.

25. Nguyen P, et al. (2008) BAT3 and SET1A form a complex with CTCFL/BORIS to modulate H3K4 histone dimethylation and gene expression. Mol Cell Biol 28(21):6720-6729.

26. Krenciute G, et al. (2013) Nuclear BAG6-UBL4A-GET4 complex mediates DNA damage signaling and cell death. J Biol Chem 288(28):20547-20557.

27. Wainright PO, Hinkle G, Sogin ML, & Stickel SK (1993) Monophyletic origins of the metazoa: an evolutionary link with fungi. Science 260(5106):340-342.

28. Baldauf SL & Palmer JD (1993) Animals and fungi are each other’s closest relatives: congruent evidence from multiple proteins. Proc Natl Acad Sci U S A 90(24):11558-11562.

29. Linding R, et al. (2003) Protein disorder prediction: implications for structural proteomics. Structure 11(11):1453-1459.

30. Xu Y, Anderson DE, & Ye Y (2016) The HECT domain ubiquitin ligase HUWE1 targets unassembled soluble proteins for degradation. Cell Discov 2:16040.

31. Hessa T, et al. (2011) Protein targeting and degradation are coupled for elimination of mislocalized proteins. Nature 475(7356):394-397.

32. Payapilly A & High S (2014) BAG6 regulates the quality control of a polytopic ERAD substrate. J Cell Sci 127(Pt 13):2898-2909.

33. Minami R, et al. (2010) BAG-6 is essential for selective elimination of defective proteasomal substrates. J Cell Biol 190(4):637-650.

34. Xu Y, Liu Y, Lee JG, & Ye Y (2013) A ubiquitin-like domain recruits an oligomeric chaperone to a retrotranslocation complex in endoplasmic reticulum-associated degradation. J Biol Chem 288(25):18068-18076.

35. Melikova MS, Kondratov KA, & Kornilova ES (2006) Two different stages of epidermal growth factor (EGF) receptor endocytosis are sensitive to free ubiquitin depletion produced by proteasome inhibitor MG132. Cell Biol Int 30(1):31-43.

36. Rodrigo-Brenni MC, Gutierrez E, & Hegde RS (2014) Cytosolic quality control of mislocalized proteins requires RNF126 recruitment to Bag6. Mol Cell 55(2):227-237.

37. Hoelz A, Debler EW, & Blobel G (2011) The structure of the nuclear pore complex. Annu Rev Biochem 80:613-643.

38. Pumroy RA & Cingolani G (2015) Diversification of importin-alpha isoforms in cellular trafficking and disease states. Biochem J 466(1):13-28.

39. Lange A, et al. (2007) Classical nuclear localization signals: definition, function, and interaction with importin alpha. J Biol Chem 282(8):5101-5105.

40. Wu YH, Shih SF, & Lin JY (2004) Ricin triggers apoptotic morphological changes through caspase-3 cleavage of BAT3. J Biol Chem 279(18):19264-19275.

41. Rouillard AD, et al. (2016) The harmonizome: a collection of processed datasets gathered to serve and mine knowledge about genes and proteins. Database (Oxford) 2016.

42. Shao S, Rodrigo-Brenni MC, Kivlen MH, & Hegde RS (2017) Mechanistic basis for a molecular triage reaction. Science 355(6322):298-302.

43. Rome ME, Chio US, Rao M, Gristick H, & Shan SO (2014) Differential gradients of interaction affinities drive efficient targeting and recycling in the GET pathway. Proc Natl Acad Sci U S A 111(46):E4929-4935.

44. Beg AA, et al. (1992) I kappa B interacts with the nuclear localization sequences of the subunits of NF-kappa B: a mechanism for cytoplasmic retention. Genes Dev 6(10):1899-1913.

45. Gu L, et al. (2011) Intermolecular masking of the HIV-1 Rev NLS by the cellular protein HIC: novel insights into the regulation of Rev nuclear import. Retrovirology 8:17.

46. Davies RG, Wagstaff KM, McLaughlin EA, Loveland KL, & Jans DA (2013) The BRCA1-binding protein BRAP2 can act as a cytoplasmic retention factor for nuclear and nuclear envelope-localizing testicular proteins. Biochim Biophys Acta 1833(12):3436-3444.

47. Wang Y, et al. (2014) GdX/UBL4A specifically stabilizes the TC45/STAT3 association and promotes dephosphorylation of STAT3 to repress tumorigenesis. Mol Cell 53(5):752-765.

48. Arhzaouy K & Ramezani-Rad M (2012) Nuclear import of UBL-domain protein Mdy2 is required for heat-induced stress response in Saccharomyces cerevisiae. PLoS One 7(12):e52956.

49. Etokebe GE, et al. (2015) Association of the FAM46A gene VNTRs and BAG6 rs3117582 SNP with non small cell lung cancer (NSCLC) in Croatian and Norwegian populations. PLoS One 10(4):e0122651.

50. Zhao J, Wang H, Hu W, & Jin Y (2014) Effect of HLA-B-associated transcript 3 polymorphisms on lung cancer risk: a meta-analysis. Med Sci Monit 20:2461-2465.

51. Chen J, Zang YS, & Xiu Q (2014) BAT3 rs1052486 and rs3117582 polymorphisms are associated with lung cancer risk: a meta-analysis. Tumour Biol 35(10):9855-9858.

52. Etokebe GE, et al. (2015) Susceptibility to large-joint osteoarthritis (hip and knee) is associated with BAG6 rs3117582 SNP and the VNTR polymorphism in the second exon of the FAM46A gene on chromosome 6. J Orthop Res 33(1):56-62.

53. Luce MJ, Akpawu AA, Tucunduva DC, Mason S, & Scott MS (2016) Extent of pre-translational regulation for the control of nucleocytoplasmic protein localization. BMC Genomics 17:472.

54. Bailey TL, et al. (2009) MEME SUITE: tools for motif discovery and searching. Nucleic Acids Res 37(Web Server issue):W202-208.

